# Dual-task effects on locomotor savings in aging

**DOI:** 10.64898/2026.05.21.726517

**Authors:** Marisa E. Mulvey, Julia T. Choi

**Affiliations:** Department of Applied Physiology and Kinesiology, University of Florida, Gainesville, FL, USA 32611

## Abstract

Healthy young and older adults completed two randomized sessions of split-belt treadmill walking, with and without a concurrent cognitive task. When the single-task session occurred first, both age groups showed savings in step length asymmetry during re-adaptation one week later. However, performing the dual-task session first reduced savings, and this order-effect was greater in older adults compared to young adults. These findings suggest that cognitive load during initial motor adaptation interferes with savings, but once stored, locomotor readaptation is resilient to dual-tasking.

## Introduction

Locomotor adaptation is the process of adjusting walking patterns in response to changing conditions^1^. Repeated exposure to the same walking perturbation, such as walking on a split-belt treadmill^2^, leads to after-effects on removal of the perturbation and savings on subsequent exposure^3-5^. Locomotor adaptation is slower when attention is divided by a concurrent cognitive task^6^, with greater motor-cognitive interference at the onset of adaptation (i.e., early adaptation) when cognitive demands are highest^7-9^. The impact of cognitive load on long-term retention of walking patterns is less clear, with one study suggesting that dual-tasking has no significant effects on savings assessed within the same session^9^.

Aging affects both the ability and strategies for learning new walking patterns^10-14^. Older adults adapt walking on a split-belt treadmill slower than young adults, even in the absence of competing cognitive tasks^15,16^. When performing a concurrent cognitive task during locomotor adaptation, older adults show further reduction in gait adaptation compared to single-task conditions^17,18^. To compensate, older adults tend to prioritize walking adaptation over cognitive task performance under dual-task conditions^14^. With repeated exposures, experienced older adults can switch more efficiently between walking patterns during locomotor adaptation^13^. While aging is often associated with memory decline^17^, one study showed that older adults can retain savings for at least one week^19^, indicating the potential for long-term locomotor memories.

Although both increased cognitive load and aging can interfere with locomotor adaptation, it remains unclear how task order affects locomotor savings in older adults. The current study investigated the effects of task order (e.g., dual → single vs. single → dual) on split-belt walking adaptation in young and older adults, with savings assessed one week later. We hypothesized that initial adaptation under higher cognitive load (dual → single) would reduce subsequent savings compared to the reverse order (single → dual). Consistent with this hypothesis, re-adaptation would be less affected by dual-tasking across age groups due to reduced attentional demands with prior experience.

## Results

Forty-one individuals participated in the study, including 19 older adults (OA: ages 55-83) and 22 younger adults (YA: ages 19-27). All participants completed two split-belt treadmill sessions (single-task and dual-task) one week apart in randomized order. Each session included baseline, adaptation (split-belt: dominant leg fast, non-dominant leg slow), and de-adaptation phases (Figure 1a). In the dual-task session, participants performed subtractions throughout walking and during seated trials before and after walking. Participants within each age group were assigned to dual-task first (OA1 and YA1) or single-task first (OA2 and YA2). Group-level participant characteristics are summarized in Table 1.

**Table 1.** Participant characteristics by group.

|  | Young adults |  |  | Older adults |  |  |
| --- | --- | --- | --- | --- | --- | --- |
|  | Dual-task first<br>**YA Group 1** | Single-task first<br>**YA Group 2** | p-value | Dual-task first<br>**OA Group 1** | Dual-task first<br>**OA Group 2** | p-value |
| Number of participants | 10 | 12 | - | 10 | 9 | - |
| Age (years) | 22.5 ± 2.4 | 21.5 ± 2.6 | 0.5 <sup>1</sup> | 73.9 ± 7.8 | 74.1 ± 9.4 | 0.7 <sup>1</sup> |
| Sex | 6 female: 4 male | 5 female: 7 male | 0.7 <sup>2</sup> | 5 female: 5 male | 7 female: 2 male | 0.3 <sup>2</sup> |
| Height (inches) | 67.2 ± 4.6 | 68.5 ± 3.9 | 0.4 <sup>1</sup> | 64.9 ± 3.2 | 64.6 ± 3.0 | 0.6 <sup>1</sup> |
| Weight (pounds) | 147 ± 27.1 | 171 ± 40.2 | 0.09 <sup>1</sup> | 150 ± 29.3 | 165 ± 20.1 | 0.2 <sup>1</sup> |
| Dominant Leg | 1 left: 9 right | 0 left: 12 right | 0.5 <sup>2</sup> | 1 left: 9 right | 2 left: 7 right | 0.6 <sup>2</sup> |
| TICS | 34.9 ± 3.8 | 36.8 ± 2.0 | 0.2 <sup>1</sup> | 36.2 ± 2.0 | 35.9 ± 1.8 | 0.8 <sup>1</sup> |
| SPPB | - | - | - | 10.6 ± 1.0 | 11.0 ± 1.0 | 0.3 <sup>1</sup> |
| Preferred gait speed (m/s) | - | - | - | 0.85 ± 0.13 | 0.87 ± 0.13 | 0.3 <sup>1</sup> |

Values are means ± SD
TICS = Telephone Interview for Cognitive Status, out of 41
SPPB = Short Physical Performance Battery, scores range from 0 to 12
<sup>1</sup> Mann-Whitney test
<sup>2</sup> Fisher's exact test

**Figure 1.**
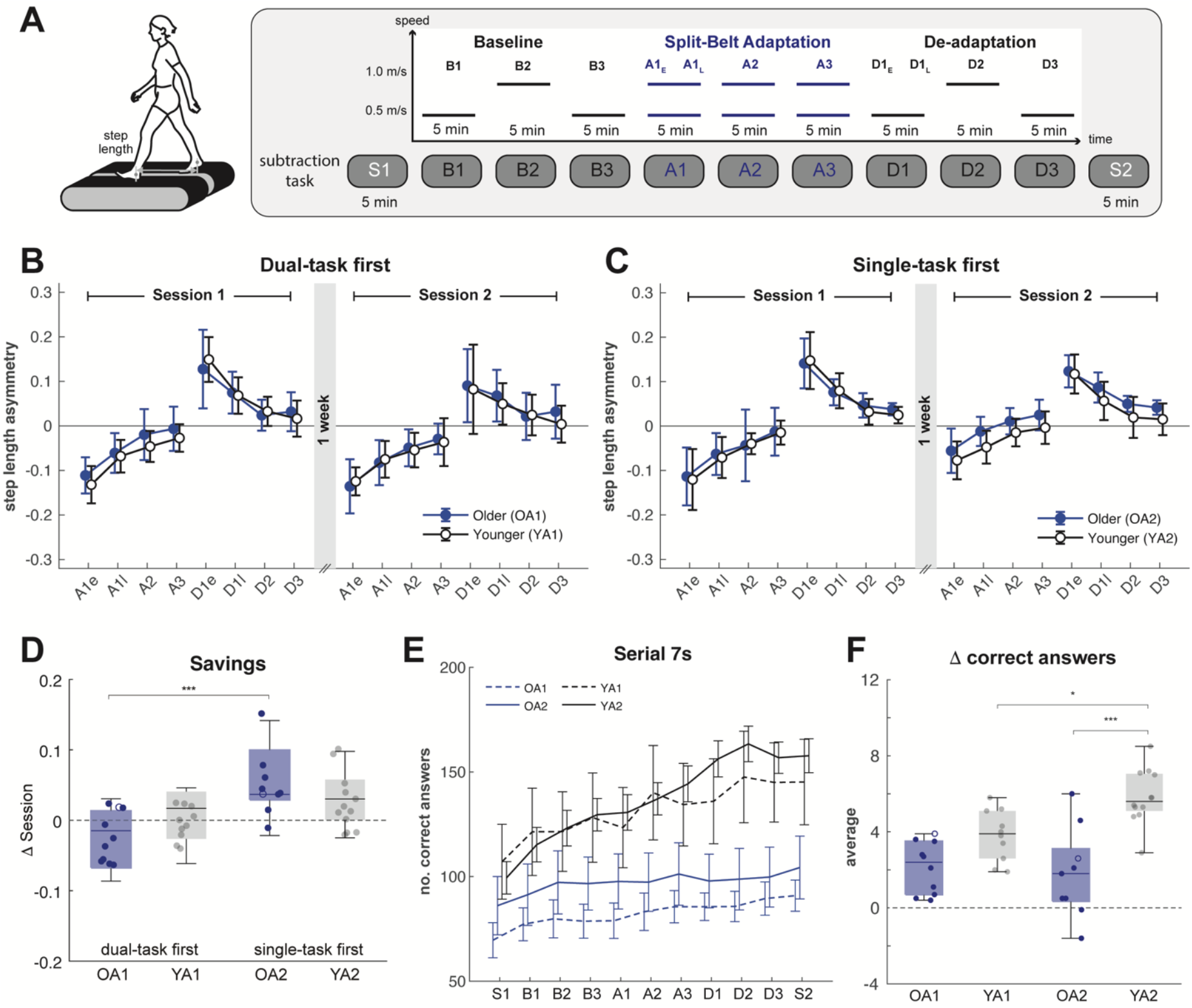
Split-belt walking adaptation. **A)** The protocol consisted of baseline, adaptation, and de-adaptation trials. Participants performed a concurrent subtraction task in the dual-task session. **B-C)** Step length asymmetry (mean ± SEM) across adaptation and de-adaptation epochs. Group average for OA1 and YA1 who completed the dual-task first (**B**), and OA2 and YA2 who completed the single-task first (**C**). Zero indicates perfect symmetry. **D)** Savings (difference between sessions) during the A1e adaptation epoch. Positive values indicate more savings. Boxes show IQR, line indicate medians, and whiskers extend to 1.5 x IQR. *** indicate post hoc comparisons with *p* < 0.001. **E)** Cognitive task performance (mean ± SEM) across trials. **F)** Change in correct answers (averaged across trials) for each participant group. * Post hoc test: *p* < 0.05. Dots represent individual participants (Fig. 1D, 1F); two middle-aged participants (< 65 years) are represented by open symbols ‘o’ to distinguish them from the rest of the sample.

All participant groups showed changes in step length asymmetry across adaptation epochs (F(1.3,47.9) = 151.8, *p* < 0.001), starting negative before approaching symmetry. The pattern was reversed during de-adaptation (F(1.3,61.8) = 165.0, *p* < 0.001).

For adaptation, there was a main effect of session (F(1,111) = 4.03, *p* < 0.05) and a significant interaction between session x task order (F(1,111) = 20.1, *p* < 0.001). There was no significant main effect of task-order (F(1,37) = 1.18 *p* = 0.3) and no significant main effect of age group (F(1,37) = 0.936, *p* = 0.3). Post hoc analysis showed that when the dual task was performed first, there were no session effects on step length asymmetry (Cohen’s *d* = -0.205, *p* = 0.6) (Figure 1b). In contrast, there was a significant session effect (Cohen’s *d* = 0.735, *p* < 0.001) when the single-task task orderwas performed first (Figure 1c).

For de-adaptation, there was a main effect of session (F(1,105) = 61.6, *p* < 0.001) but no significant interaction of session x task order (F(1,105) = 1.04, *p* = 0.03). There were also no significant main effects of task-order (F(1,37) = 0.174, *p* = 0.7) nor age group (F(1,37) = 1.58, *p* = 0.2).

Savings, calculated as the difference between sessions, showed a significant effect of task order (F(1, 37) = 13.3, *p* < 0.001) (Figure 1d). There was no significant main effect of age group (F(1,37) = 0.291, *p* = 0.6) and no interaction of age x task order interaction (F(1,37) = 2.1, *p* = 0.2). For A1e, the effect size for task-order was large (Cohen’s *d* = -1.19, *p* < 0.001) with a greater effect in older adults (OA1 vs. OA2: Cohen’s *d* = -1.60, *p* = 0.007) compared to young adults (YA1 vs. YA2: Cohen’s *d* = -0.692, *p* = 0.4). A second analysis on savings across adaptation epochs similarly showed a significant effect of task order (F(1, 37) = 20.1, *p* < 0.001), no significant effect of age group (F(1,37) = 0.03, *p* = 0.8), no significant interaction of age x task order interaction (F(1,37) = 3.72, *p* = 0.06), and no significant effect of epoch on savings (F(2.0, 73.8) = 2.68, *p* = 0.08).

Older adults had fewer correct answers and showed less improvement (Δ correct answers) across trials than young adults in the subtraction task (Figure 1e). There was a significant effect of age (F(1,37) = 30.1, *p* < 0.001) and an age x order group interaction (F(1,37) = 5.0, *p* = 0.03) (Figure 1f). Young adults who performed the single-task task order first improved more than those who performed it second (YA1 vs. YA2; Cohen’s *d* = -0.15; *p* = 0.03). However, older adults who performed the single-task task order first improved less than young adults in the same order group (OA2 vs. YA2: Cohen’s *d* = - 0.30, *p* < 0.001), and there were no differences between the two groups of older adults (OA1 vs. OA2: Cohen’s *d* = 0.02, *p* = 0.7).

## Discussion

Our findings provide novel insights into locomotor learning in aging: 1) older adults demonstrated locomotor savings after a 1-week interval; and 2) savings were reduced when the initial adaptation included a cognitive task, suggesting that dividing attention interfered with consolidation of locomotor memory. Older adults exhibited greater locomotor savings following single-task adaptation, while younger adults showed more improvement in cognitive task performance.

Savings reflects the ability to retrieve learned motor strategies or memories^20,21^. We found that locomotor savings were absent following dual-task adaptation, contrasting with a previous report showing savings after both single- and dual-task split-belt adaptation^9^. This difference may be due to the time interval between initial adaptation and re-adaptation. The prior study assessed immediate savings within the same session, which may rely on short-term processes. In contrast, 1-week savings likely involves long-term memory consolidation^22^.

Outside the split-belt literature, visuomotor studies have shown that attentional context can influence savings^23^. To our knowledge, this is the first evidence of a directional effect in locomotor adaptation (dual-single vs. single-dual). This pattern aligns with sensorimotor adaptation models proposing that savings primarily reflect the recall of an action-selection strategy, rather than error-based adaptation^24^. Dual-tasking during the initial exposure may disrupt this strategic component that supports savings, which could explain why task order mattered in this study.

Given that dual→single reduced savings relative to single→dual, it is reasonable to infer that dual→single would also show reduced savings compared to a single→single group. Our study did not include a single-single group. However, published data on split-belt locomotor adaptation in middle-aged adults provide single-single effect sizes for comparison.^9^ Previously reported savings in step length asymmetry (Rossi et al. 2021 Figure 4B: mean ± SE = 0.039 ± 0.02) are comparable to the A1e savings observed in our single-dual groups (OA: 0.058 ± 0.02 and YA: 0.043 ± 0.02), suggesting that single-single and single-dual sessions have similar magnitude of savings.

Since rehabilitation sessions are often spaced over days or weeks^25-27^, the presence of 1-week locomotor savings is clinically important. Our findings suggest that starting with single-task training may enhance motor savings when followed by progressive introduction of dual-task challenges to promote real-world transfer. Understanding how cognitive processes interacts with locomotor savings is critical for designing effective interventions. Older participants in this study likely represent a high-functioning segment of the aging population, as they had normal cognition (TICS = 36, corresponding to MMSE = 30)^28^ and gait speed within normative range^29^. All participants tolerated a 0.5 m/s belt-speed difference and demonstrated typical locomotor adaptation capacity. In contrast, individuals with greater mobility or cognitive impairments may exhibit reduced locomotor adaptation^30^. Whether such impairments increase dual-task costs or limit locomotor savings remains to be determined.

A limitation of this brief report is that we focused primarily on step length asymmetry, a commonly reported measure in split-belt adaptation studies, to allow comparison with prior work^9,13,14,17^. Future studies should examine other gait parameters (e.g., double support time) and consider the influence of different cognitive tasks on locomotor adaptation and savings in older adults. Additionally, testing longer time intervals would clarify the durability of locomotor savings and its clinical implications.

## Methods

### Participants

Healthy young adults (YA; ages 19-27) and typical aging older adults (OA; ages 55-83) were included in this study. Individuals with neurological disorders were excluded. Cognitive status was pre-screened using the Telephone Interview for Cognitive Status (TICS), which strongly correlates with the Mini-Mental State Examination.^28,31^ Participants with possible dementia (TICS ≤ 29)^32^ were excluded. Older adults completed the Short Physical Performance Battery (SPPB), and their preferred gait speed was calculated from two trials of the 4-m walk. Dominant leg was determined using the Waterloo footedness questionnaire^33^. None of the participants had previous split-belt treadmill experience. All participants provided written informed consent prior to participation. The protocol was approved by the Institutional Review Board of the University of Florida (IRB201902465) and according to the Declaration of Helsinki.

### Experimental Design

Split-belt walking was performed under single-task and dual-task conditions over two sessions, separated by one week, and assigned in random order (Figure1a). Participants were divided into four groups based on task order and age: OA1 (n = 10) and YA1 (n = 10) completed the dual task first, while OA2 (n = 9) and YA2 (n = 12) completed the single task first (Table 1).

Participants walked on a split-belt treadmill with independent belt speed control (Bertec, Columbus, OH, USA). Each session began with three 5-minute tied-belt trials at 0.5 m/s, 1.0 m/s, and 0.5 m/s (baseline: B1, B2, and B3, respectively). This was followed by three 5-minute split-belt trials (adaptation: A1, A2, and A3, respectively), where the dominant leg (“fast leg”) moved at 1.0 m/s and the non-dominant leg (“slow leg”) at 0.5 m/s. The session concluded with three 5-minute tied-belt trials at 0.5 m/s, 1.0 m/s, and 0.5 m/s (de-adaptation: D1, D2, and D3, respectively). The treadmill was stopped briefly (about 30 seconds) after each 5-minute trial, and participants remained standing on the treadmill during this period.

In the dual-task condition, participants performed a subtraction task during walking and while sitting before (S1) and after (S2) walking. Serial subtractions by sevens was used because this task reliably induces cognitive-motor interference during gait in older adults.^34,35^ Participants were instructed to “count backwards by seven as quickly and as accurately as possible” from a random number. Each iteration involved 14 subtractions and was repeated with a new starting number throughout the 5-minute trials.

### Data Collection

Reflective markers were placed on the left and right foot (fifth metatarsals) and tracked at 100 Hz using 8 Miqus cameras (Qualysis, Gothenburg, Sweden). A Qualisys sync unit was used to synchronize motion capture and force data. Ground reaction forces (GRFs) were recorded from the force plates under each treadmill belt at 1000 Hz.

### Data Analysis

Data analysis was performed in MATLAB (version 2023b, MathWorks, Natick, MA, USA). Force data was low-pass filtered using a 3^rd^ order Butterworth with 15 Hz cut-off, applied in the forward and backward directions for zero-phase filtering. Heel-strike events were identified from the vertical GRFs using a threshold of 0.08 x body weight,^36^ then visually inspected and manually corrected for any erroneous or missing events. Step length was defined as the anterior-posterior distance between the left and right markers at heel-strike.^36^ Step length asymmetry was calculated as the normalized difference:

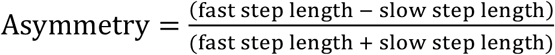

Mean step length asymmetry was calculated for the following epochs: the first 30 strides of A1 (A1e), strides 31 through the end of A1 (A1l), all strides from A2, and all strides from A3. A1 was split into A1e and Al1 to capture early and late adaptation, respectively. A1e includes the period identified in prior work as having the largest reduction in step length asymmetry, while A1l reflects later changes in adaptation.^21^ The same bins were applied to de-adaptation. Savings was calculated as the difference in step length asymmetry between sessions for each adaptation epoch, where positive values indicated greater savings.

### Cognitive Task Performance

Performance in the serial 7s subtraction task was based on the number of correct answers (total responses minus errors) for each 5-minute interval. To measure changes in performance, we calculated the difference in correct answers from the previous trial (Δ correct answers). This measure accounts for learning across trials and is relative to each participant’s own baseline performance.

### Statistical Analysis

Statistical analyses were conducted in JASP Version 0.16.2 (JASP Team, 2016, jasp-stats.org). Changes in step length asymmetry were analyzed separately for adaptation (A1e, A1l, A2, A3) and de-adaptation (D1e, D1l, D2, D3) epochs. A mixed-measures ANOVA was performed with session (session 1 vs. session 2) and epoch as within-subject factors, and age group (OA vs. YA) and task-order (dual-task first vs. single-task first) as between-subject factors. Mauchly’s test indicated that the assumption of sphericity was not met for epoch (χ^2^ = 65.3, *p* < 0.001); degrees of freedom were corrected using Greenhouse-Geisser. Post-hoc comparisons were adjusted using the Holm correction for multiple comparisons, and effect sizes (Cohen’s *d*) were computed. A *p*-value less than 0.05 was considered statistically significant.

A1e savings was used as the primary metric. Savings was analyzed using a two-way ANOVA with task-order and age group as a between-subject factors. Effect sizes (Cohen’s *d*) were computed for the main effect of task-order, and for task-order effects within each age group. A secondary analysis examined whether savings differed across adaptation epochs using a mixed-measures ANOVA with task-order and age group as a between-subject factors, and epochs (A1e, A1l, A2, A3) as a within-subject factor.

For the subtraction task, we analyzed the change in correct answers (Δ correct answers) using a mixed-measures ANOVA with trial as the within-subject factor and age group and task order as between-subject factors. The number of correct answers during the first sitting trial (S1) was included as a covariate to account for initial performance.

### Sample Size

The study was planned to include 40 participants (n = 10 per group). This sample size was based on previous studies of age-related differences in dual-task split-belt adaptation, which included 10 to 14 participants per group.^9,14,18^ Based on the planned sample size and a two-sided α = 0.05, the ANOVA was powered to detect moderate-to-large effects (f = 0.40). For main effects, this corresponds to large effects (Cohen’s *d* = 0.8). The final sample size was comparable and provided similar sensitivity.

## Clinical trial number

not applicable

## Data Availability

The data that support the findings of this study are available upon reasonable request from the corresponding author.

## Author contribution

MM and JC conceived and designed the study. MM performed the experiments and analyzed the data. MM drafted the manuscript. JC created the figures and revised the manuscript. MM and JC approved the final version of manuscript.

## Acknowledgements

We sincerely thank Dr. Muxuan Liang for valuable input on statistical considerations.

